# The structural diversity of telomeres and centromeres across mouse subspecies revealed by complete assemblies

**DOI:** 10.1101/2024.10.24.619615

**Authors:** Bailey Francis, Landen Gozashti, Kevin Costello, Takaoki Kasahara, Olivia S. Harringmeyer, Jingtao Lilue, Mohab Helmy, Tadafumi Kato, Anne Czechanski, Michael A. Quail, Iraad Bonner, Emma Dawson, Anne Ferguson-Smith, Laura Reinholdt, David J. Adams, Thomas M. Keane

## Abstract

It is over twenty years since the publication of the C57BL/6J mouse reference genome, which has been a key catalyst for understanding mammalian disease biology. However, the mouse reference genome still lacks telomeres and centromeres, contains 281 chromosomal sequence gaps, and only partially represents many biomedically relevant loci. We present the first T2T mouse genomes for two key inbred strains, C57BL/6J and CAST/EiJ. These T2T genomes reveal significant variability in telomere and centromere sizes and structural organisation. We add an additional 213 Mbp of novel sequence to the reference genome containing 517 protein-coding genes. We examined two important but incomplete loci in the mouse genome - the pseudoautosomal region (PAR) on the sex chromosomes and KRAB zinc finger proteins (KZFPs) loci. We identified distant locations of the PAR boundary, different copy number and sizes of segmental duplications, and a multitude of amino acid substitution mutations in PAR genes.

## Introduction

Mice have been used for a hundred years to model human disease leading to key discoveries such as the role of the H2/MHC locus in immunity ^1^, the discovery of oncogenes and tumour suppressors ^2^, and the development of induced pluripotent stem cells (iPSCs) ^3^. Research using mice has provided researchers with a valuable tool to study disease mechanisms, develop treatments, and discover the genetic basis for physiological processes.

In 2002, the generation and assembly of the first mouse genome for the C57BL/6J strain represented a major milestone for mouse genetics^4^. The mouse karyotype consists of 19 pairs of telocentric autosomes and X chromosome, having no obvious short arm apart from the Y chromosome, which is acrocentric ^5,6^. Telocentric chromosomes are challenging to fully assemble due to large satellite arrays near the centromere and telomere end. The current mouse genome (GRCm39) remains incomplete due to 281 gaps distributed across every chromosome, a partial set of telomeres, and no centromeres. Telomeres and centromeres are critical structural components of chromosomes, each playing a unique role in maintaining chromosomal stability and integrity. High throughput sequencing of ultra long DNA fragments (>100Kbp) represents a unique opportunity to assemble fully complete Telomere-to-Telomere (T2T) chromosomes, recently demonstrated by the first human T2T genome ^7^.

In this study, we have used single molecule ultra-long sequencing to produce the first T2T mouse genome for two inbred strains that represent two subspecies of *Mus musculus*; C57BL/6J and CAST/EiJ. We show how these T2T reference genomes are more complete than the current mouse genome (GRCm39), notably adding complete telomeres and centromeres for all autosomes, sequence across current gaps in GRCm39, and completing important loci such as the pseudoautosomal region (PAR) on CAST/EiJ chromosome X and two KRAB zinc finger proteins (KZFPs) clusters.

## Results

We obtained DNA from embryonic stem cells derived from a CAST/EiJ x C57BL/6J F1 male embryo. Sequencing was performed using a combination of Pacbio HiFi (42x coverage) and Oxford Nanopore ultra long sequencing (52x coverage, 22x was >100kbp read length). We employed a trio-based genome assembly approach using parental short reads to assign each long read to its parental haplotype (see methods). Genome assemblies were generated using both Verkko ^8^ and Hifiasm ^9^. We produced six distinct assemblies using both assemblers, and applied a set of quality control measures to select the single best base assembly. We assessed the k-mer completeness to compute an overall quality value (QV) score for each assembly. Haplotype separation was evaluated by comparing k-mer spectrums of the two haploid assemblies against k-mer spectrums from parental strains sequencing data, and comparing to a combined reference genome of GRCm39 and a previous Pacbio long read CAST/EiJ assembly (GCA_921999005.2). We searched for mouse canonical telomeric repeats (TTAGGG) at the end of the chromosome contigs and used telomere presence as a marker for complete chromosome ends. We selected the best initial base assembly for each strain by comparing and ranking various assembly quality metrics (Supplementary Table 1). For both strains, we selected a Verkko assembly as the base, which was then improved by a round of curation (see methods), and a round of polishing which improved the base accuracy.

Several chromosomes still did not end in telomeric sequence at the centromeric end. We identified the missing telomere sequences by searching for the mouse canonical telomere repeat in the unplaced contigs. Human studies noted that large satellite arrays tend to have more similarity within a given chromosome array than between different chromosomes ^10^. We applied this method in combination with long range Hi-C mate pairs from the parental strains to identify the chromosome scaffold of greatest similarity and support from Hi-C mate pairs (see methods). This allowed us to assign all remaining telocentric sequences to a chromosome.

Table 1 provides an overview of the final C57BL/6J and CAST/EiJ T2T assemblies. The autosomes’ ungapped length for both assemblies is consistently longer than GRCm39, adding 208 (C57BL/6J) and 247 Mbp (CAST/EiJ) of additional sequence. Base accuracy was higher in the T2T C57BL/6J than GRCm39 (QV 54.9 vs. QV 54.4, respectively), and slightly lower for CAST/EiJ (QV 48.2). Mapping of parental short reads to GRCm39 and T2T C57BL/6J shows an increase of 0.7% mapped read pairs and 0.22% correctly paired reads in the C57BL/6J T2T genome (Supplementary Table 2). Significant deviations in mapped reads are observed on chromosomes 2 and 9 due to the presence of short satellite-like sequences within two regions of these chromosomes.

To compare the structural accuracy of the C57BL/6J T2T assembly and the GRCm39 reference genome, we used Pacbio reads from the C57BL/6J ‘Eve’ mouse to call structural variants ^11^. 11% fewer SVs were predicted from the T2T genome compared to GRCm39 - insertions (4,147 vs 4,496), deletions (3,192 vs 3,295), duplications (0 vs 3), and inversions (660 vs 654) (Supplementary Table 3).

### Chromosome Structure and annotation

Figure 1 provides a synteny comparison of the GRCm39, T2T C57BL/6J, and the T2T CAST/EiJ genomes highlighting the presence of telomeres and centromeric sequences projecting from the ends of the T2T mouse genomes. Gaps in GRCm39 that have been filled and expansions are visible, and the presence of large scale multi megabase inversions between C57BL/6J and CAST/EiJ strains. Telomeres and centromeres are present on the ends of each T2T genome, whereas telomeres are only present on five autosomal chromosomes of GRCm39 on the non-centromeric ends. The total novel sequence in the T2T genomes compared to GRCm39 is 213.2 Mbp and 252.1 Mbp for C57BL/6J and CAST/EiJ, respectively, and the vast majority of this consists of common repeats such as satellites and transposons (Fig. 1C). Total satellite sequence has been increased by over 3,000% in both strains compared to GRCm39, along with substantial increases in all common repeat classes except LINEs in CAST/EiJ (Fig. 1B).

**Figure 1.**
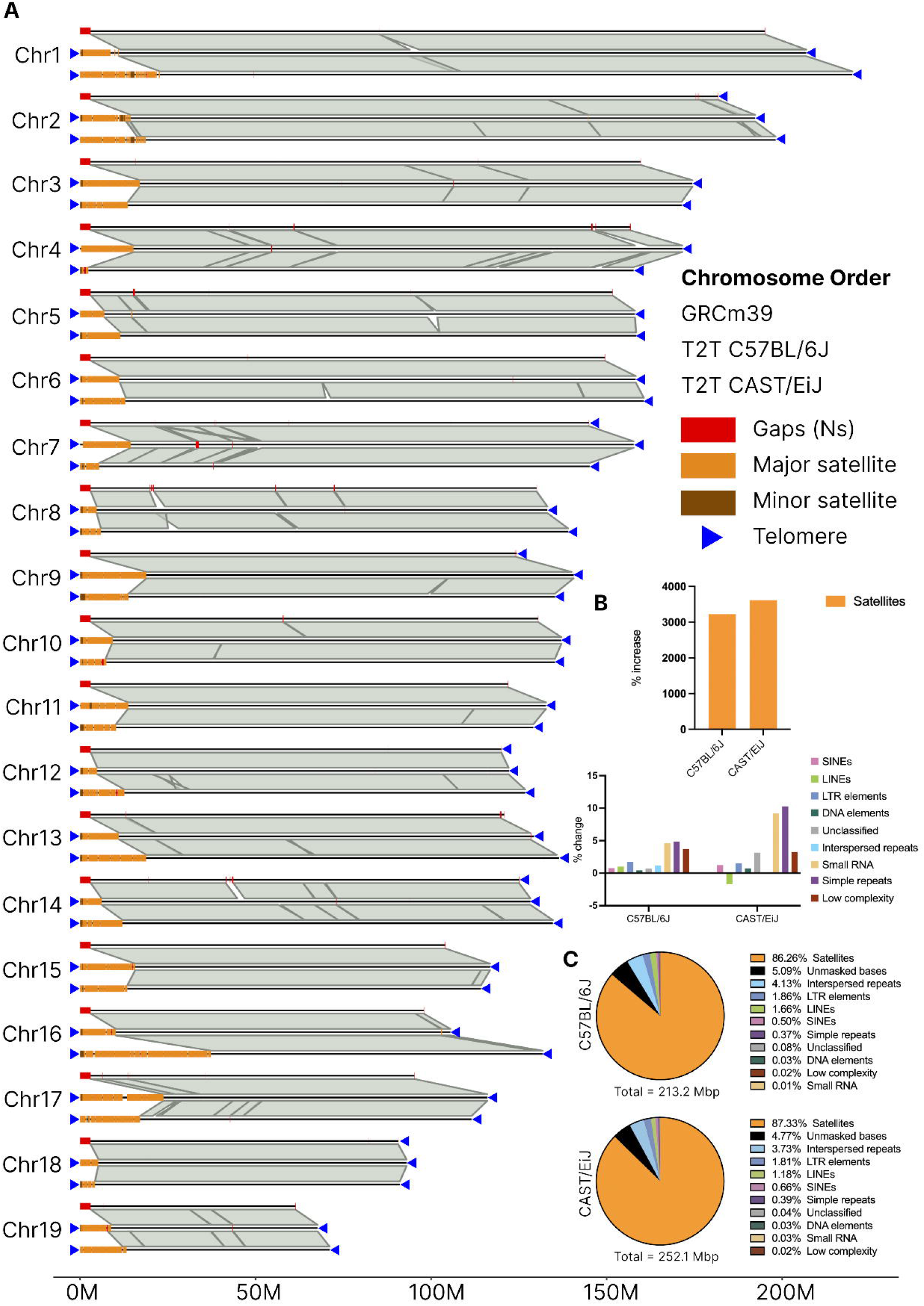
Chromosome scale synteny comparison of the mouse T2T genomes and GRCm39. A) Three-way synteny comparison of GRCm39 (top), T2T C57BL/6J (middle), and T2T CAST/EiJ (bottom) for all chromosomes. Gaps, centromeres (major and minor mouse satellites), and telomeres (where present) are annotated on each chromosome. B) Increases in satellite sequence and common repeat classes in the T2T genomes relative to GRCm39. C) Categorisation of repeat content of novel sequence in the T2T genomes with respect to GRCm39 (centromeres, telomeres, and gap-filling sequences).

Gene annotation was carried out using RNA-Seq from brain, liver and various cell types for C57BL/6J and liver, brain, olfactory, spleen, testis, B-cell and T-cell for CAST/EiJ (Supplementary Table 4). Total protein coding gene counts are comparable to GRCm39 (20,670 in GRCm39, 21,423 in T2T C57BL/6J, and 21,440 for T2T CAST/EiJ). We also performed gene liftover from GRCm39 to the new T2T genomes with Liftoff, which annotated 21,469 and 21,490 for C57BL/6J and CAST/EiJ, respectively.

To identify potential novel genes, we extracted genes from the BRAKER annotation that exhibited no overlap with any gene from the Liftoff annotation. Considering those with at least 3 exons and >200 bp of coding sequence, we identified 225 and 355 novel genes in C57BL/6J and CAST/EiJ, respectively (Supplementary Table 5). For C57BL/6J, novel gene size varied from 552 bp to 73,352 bp and contained between 3 to 17 exons, with a corresponding total coding sequence lengths ranging between 201 bp and 9,288 bp. In CAST/EiJ, gene size varied from 398 to 74,364 bp and contained between 3 and 25 exons, with a total coding sequence between 201 and 9,321 bp. These novel genes received significant BLAST hits to known proteins such as zinc finger proteins.

The Liftoff annotation also identified several genes from GRCm39 that have an increased copy number in the T2T genomes (Supplementary Table 6). In total, we found 201 and 247 genes with an increased copy number with respect to GRCm39 in C57BL/6J and CAST/EiJ, respectively. In both strains, *Gapdhrt2* exhibited the highest extra copy number with 31 additional copies in C57BL/6J and 35 additional copies in CAST/EiJ. Genes with increased copy number fell into a diverse range of functional categories such as immune-associated proteins, transcription factors and signal transduction proteins. Furthermore, we identified strain-specific differences in genes with increased copy number. Notable differences include genes such as *Potefam3a* and *Sp140l1*, which displayed differences of up to 22 copies between the T2T genomes.

### Telomere and centromere structure

Telomeres and centromeres are essential for chromosome integrity, maintenance, and segregation during cell division. Telomeres are repetitive nucleotide sequences at the ends of chromosomes that act as protective caps, preventing the ends of chromosomes from being recognized as DNA damage. The centromere is essential for mitotic spindle capture and checkpoint control, sister chromatid cohesion and release, and cytokinesis. GRCm39 has very limited representation of telomeres and centromeres. Historically, these sequences have been notoriously hard to resolve due to their highly repetitive nature, and as a result, are currently represented as gaps within the current mouse reference genome. These T2T mouse genomes dramatically improve the representation of these regions, allowing us to interrogate the architecture and function of mouse telomeres and centromere sequences.

In GRCm39, the canonical telomere repeat (TTAGGG)n is observed in only 6 out of 42 chromosome ends, resulting in a total telomere length of around 8 Kbp. In contrast, the T2T genomes possess full representations of telomeric sequences on all chromosome ends, amounting to a total telomere length that is around 50 times larger than GRCm39 (492 Kbp and 404Kbp in C57BL/6J and CAST/EiJ, respectively) (Supplementary Table 7). The total telomere length that we observe in the mouse chromosomes is approximately four times longer than those of the recently published human T2T assembly (121.1 Kbp) ^7^. It is well characterised that mice possess longer telomeres than humans due to higher telomerase activity ^12,13^. Figure 2C displays a comparative analysis of telomere lengths at both chromosome ends. Median telomere lengths at both the telomere end and centromere end of the chromosome are predominantly clustered around 10 Kbp for both strains. However, both C57BL/6J and CAST/EiJ exhibited a wider range of telomere lengths at the centromere end. When comparing across strains, CAST/EiJ shows a tighter distribution of telomere lengths, with less variation between the centromere and telomere ends when compared to C57BL/6J. Additionally, the longest telomeres at both chromosome ends were observed in C57BL/6J, with a maximum telomere length of 23.1 Kbp and 17.3 Kbp in C57BL/6J and CAST/EiJ respectively, both substantially lower than the BALB/cJ inbred strain ^14^. These findings are consistent with previous experiments that show CAST/EiJ to display both shorter and less variable telomere lengths than C57BL/6J ^15^.

**Figure 2.**
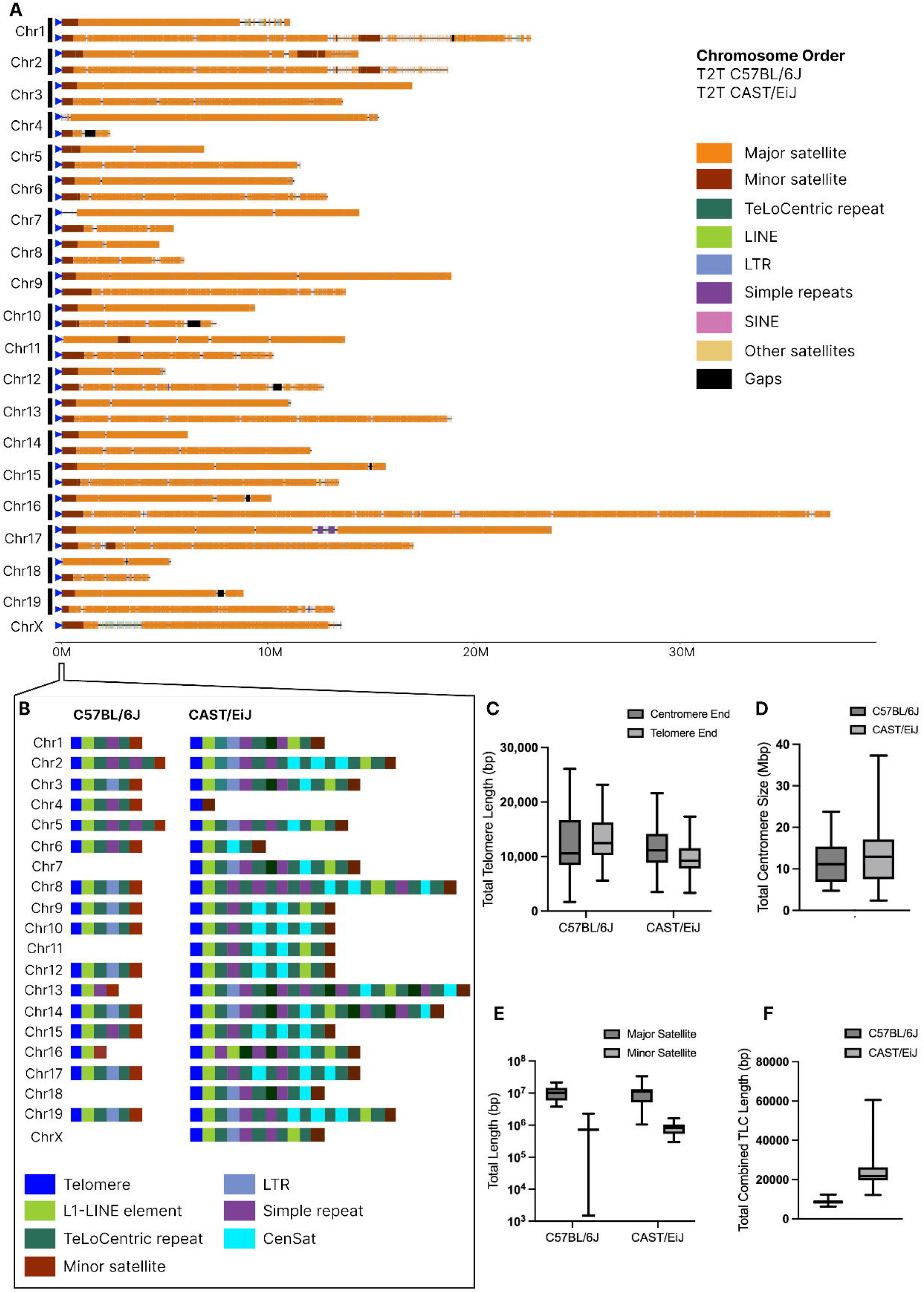
Centromere and telomere comparison of the T2T genomes. A) Comparison of centromere chromosome ends between T2T C57BL/6J and CAST/EiJ. On each chromosome, RepeatMasker repeat classes are annotated. B) Structure of the subtelomeric region (centromere side) of C57BL/6J and CAST/EiJ. C) Size distribution for canonical telomere repeat on each chromosome (TTAGGGn) (D) Centromere size distribution for T2T C57BL/6J and CAST/EiJ. E) Size distribution of length of major and minor centromere satellite sequences per chromosome. F) Size distribution of the TLC subtelomeric repeat per chromosome.

Most human chromosomes are metacentric having their centromere located in the middle of the chromosome, whereas mouse chromosomes are telocentric, with their centromeres being located at the very end of the chromosome with as little as 2 Kbp of sequence to the telomere ^16^. Figure 2A shows the location of the mouse centromeres in both strains. As expected, mouse centromeres are located directly next to the telomere highlighting that both mouse strains have telocentric chromosomes. Mouse centromeres are composed of two predominant classes of satellite DNA sequences. The most abundant class is the mouse major satellite, a 234-bp repeat monomer that has been previously reported to account for around 6-10% of the mouse genome ^17,18^. In GRCm39, major satellite sequences only occupy 99.6 Kbp of the placed chromosomes with a median total length per chromosome of 1.8 Kbp, and in the C57BL/6J and CAST/EiJ T2T genomes, they account for 210.5 Mbp and 239.7 Mbp of the placed chromosomes (12.5% and 11.8% of the genome). The minor satellite is the other predominant class of satellite DNA in mouse centromeres, an AT-rich 120-bp repeat monomer that has been previously reported to occupy 0.3-1 Mbp per chromosome ^19–21^. They are completely absent in the current mouse reference genome and are 8.6 Mbp and 12.7 Mbp total length in the C57BL/6J and CAST/EiJ T2T genomes, respectively. Comparative analysis of total centromere size revealed significant variability between mouse strains. The median total centromere size for C57BL/6J (11.1 Mbp) was smaller than in CAST/EiJ (12.9 Mbp). Additionally, the maximum centromere size observed in CAST/EiJ (36.2 Mbp, Chromosome 16) was significantly higher than C57BL/6J (23.7 Mbp, Chromosome 17) (Figure 2A, Figure 2D, Supplementary Table 8). The distribution of centromere lengths is broader in CAST/EiJ than in C57BL/6J, (interquartile range 15 Mbp for CAST/EiJ compared to 10 Mbp for C57BL/6J), and the overall range extending from around 5 to 35 Mbp in CAST/EiJ versus 5 to 25 Mbp in C57BL/6J.

C57BL/6J telocentric chromosome ends exhibit a distinct repeat organisation that is shared across chromosomes ^16,22^. These telocentric regions are characterised by stretches of the mouse canonical telomeric repeat (TTAGGG)n, followed by subtelomeric regions composed of high density repeat sequences. At the start of these subtelomeric regions, C57BL/6J exhibits a highly conserved L1-LINE element from the L1-MdA2 family. Following on from this LINE element, previous studies have described repeat arrays of mouse TeLoCentric repeat (TLC) monomers ^16,22^. These TLC arrays have been shown to be punctuated by both LTR elements and simple repeats. Finally, it has been shown that these telocentric transition sequences terminate in mouse minor satellite arrays which also denote the beginning of the mouse centromeric satellite arrays ^16,22^.

We utilised RepeatMasker and BLAST searches (see methods) to characterise the telocentric transition sequence structure. Concordant with previous findings, we found a highly conserved L1-LINE element in 16 out of 19 of C57BL/6J telocentric chromosome ends. However, the L1-element identified in the chromosome ends is a member of the L1-MdA3 family (rather than the L1-MdA2 family observed in earlier studies). Also consistent with the proposed model of C57BL/6J chromosome ends, we find TLC arrays immediately downstream of this L1-LINE element in 14 out of 16 chromosomes. The TLC arrays in the telocentric regions follow three distinct structural patterns (Figure 2B). In the first pattern, which is observed in eight chromosomes, the C57BL/6J TLC arrays are punctuated by a conserved LTR element (RLTR17B_Mm). In the remaining two patterns, the TLC arrays are instead punctuated by AT-rich simple repeats. Of the six TLC arrays that are punctuated by simple repeats, four are punctuated by a single simple repeat, whereas the remaining two are punctuated by two simple repeats. Of the 4 TLC arrays with a single simple repeat, three have the same basic structure (TATA)n -> (CATACT)n -> (TATA)n, whereas the final one is composed of (CATACT)n. The two TLC arrays with two simple repeats have the same conserved structure with the first repeat being (ACATAGTAT)n and the second repeat being (TATATGAG)n. As expected, 16 of 19 subtelomeric sequences end in minor satellite arrays.

Notable exceptions are Chromosome 7 and 11 in C57BL/6J, which do not terminate in the expected model for telocentric chromosome ends. Adjacent to the telomere, there are various repetitive elements such as SINEs, LINEs and LTRs with no clear pattern. In chromosome 11, the first centromeric satellites occur at roughly 83 Kbp with an array of major satellite sequences. The first instance of the minor satellite in this chromosome occurs at roughly 2.7 Mbp with the same L1-LINE -> TLC -> minor satellite motif observed in other chromosomes. Chromosome 7 transitions from the telomere to various LINE and LTR elements, with the first centromeric satellites appearing at roughly 750 Kbp. The first instance of minor satellite sequences in this chromosome occurs at roughly 37 Mbp.

The CAST/EiJ telocentric chromosome end structures are highly heterogeneous with no clear shared repeat organisation (Fig. 2B). Instead, CAST/EiJ appears to exhibit a set of distinct repeat motifs that appear in highly variable higher order confirmations.

However, the CAST/EiJ assembly reveals repeat motifs that are not observed in C57BL/6J telocentric chromosome ends (Fig. 2B). Firstly, centromeric satellite repeats (CenSat) repeats are observed in 18 out of 20 CAST/EiJ chromosomes, whereas this repeat is totally absent in C57BL/6J chromosome ends. These CenSat repeats are always found within TLC/CenSat repeat arrays in the CAST/EiJ chromosomes. Secondly, repeat arrays involving LTR/TLC/Simple repeats appear to be more complex in CAST/EiJ with 12 out of 20 CAST/EiJ chromosomes showing an expansion in these repeat arrays when compared to C57BL/6J. Finally, 14 out of 20 chromosomes in CAST/EiJ exhibit a second L1-LINE repeat. This L1-LINE repeat is always the same across CAST/EiJ chromosomes (L1MdGf_II) and appears to be a part of a CAST-specific repeat motif that leads into the centromere in 11 of these chromosomes: L1-LINE -> TLC -> minor satellite.

We found differences in the amount of TLC repeats between the strain assemblies. In the C57BL/6J assembly, the total amount of TLC repeat is highly conserved across chromosomes ranging from 6.2 to 12.3 Kbp. Conversely, the amount of TLC in CAST/EiJ is highly variable ranging from 12.1 to 60.5 Kbp. Additionally, the median combined TLC length is significantly higher in CAST/EiJ. These results are also concordant with previous experimental results showing significantly larger amounts of the TLC repeats in CAST/EiJ when compared to C57BL/6J ^16^.

### Completing the mouse reference genome

Despite successive efforts to fully sequence the mouse genome^11,23,24^, the GRCm39 reference genome contains approximately 87 autosomal sequence gaps estimated to be 5.5 Mbp. Gap-filling sequences were identified in the C57BL/6J T2T assembly by mapping sequences that flank gaps in GRCm39. The T2T C57BL/6J assembly completely spans 80 of these gaps (92%) and has partial closure of the remaining 7 gaps, introducing roughly 12.7 Mbp of novel sequence to the mouse genome (Figure 3A, Supplementary Table 9). By lifting over gene annotations from GRCm39 to the T2T C57BL/6J assembly, we observe a total of 301 protein-coding genes within novel gap-filling sequences (Figure 3B). Functional characterisation of these genes with PANTHER revealed that the majority of these genes fall into the transmembrane signal receptor category. Other notable categories included gene-specific transcriptional regulators, transfer/carrier proteins, and chromatin-associated proteins.

**Figure 3.**
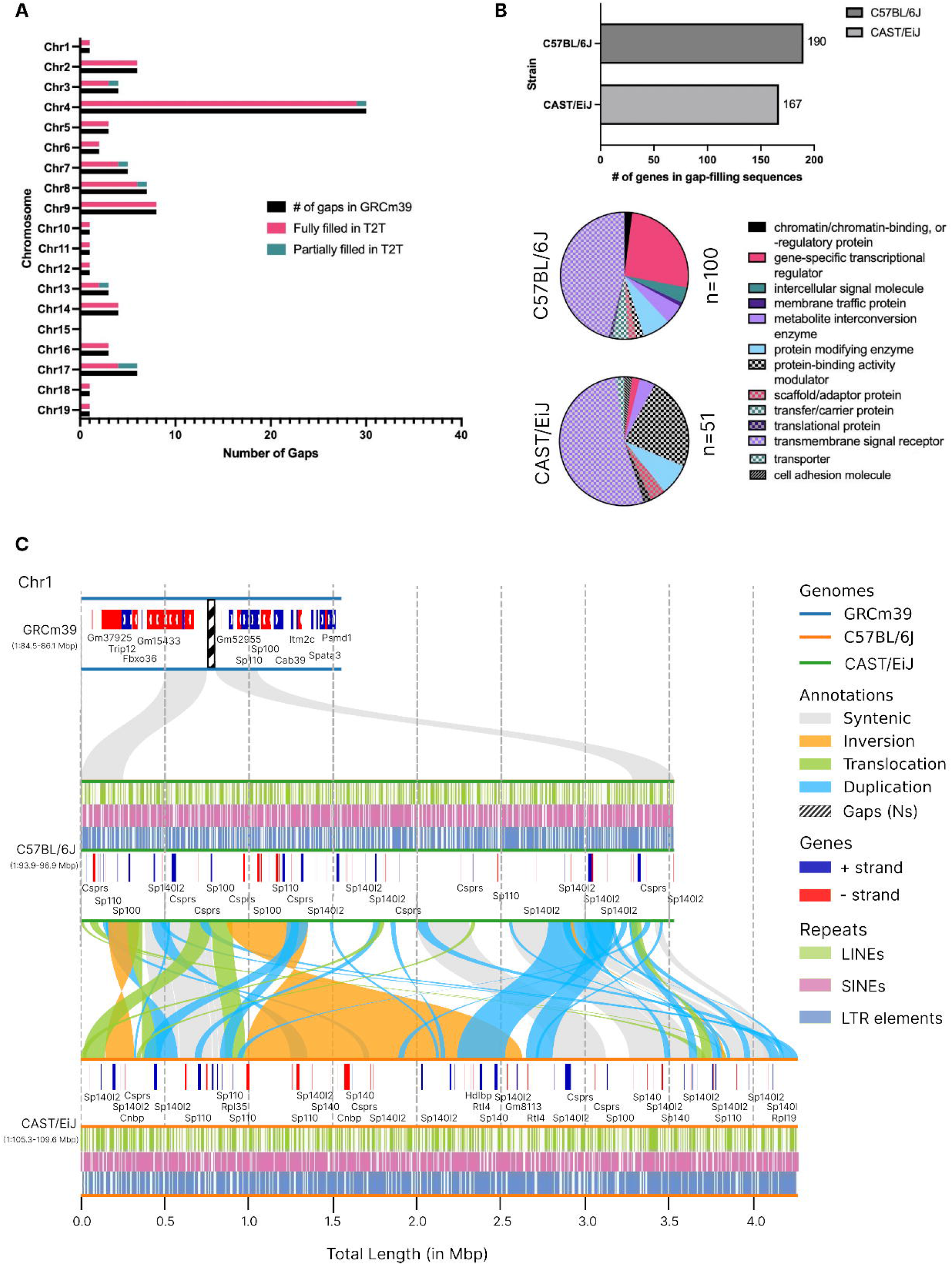
Overview, status, and content of the gap regions of GRCm39 in the T2T C57BL/6J genome. A) Chromosome breakdown of the status of the gaps GRCm39 in the T2T C57BL/6J. B) Number of protein coding genes predicted and the protein classes in the novel gap-filling sequence in the T2T C57BL/6J. Genes with no PANTHER protein class were omitted from pie charts. C) Example of a completely filled gap on Chr1 that expanded from 50 kbp in GRCm39 to 3.03 Mbp in T2T C57BL/6J. A synteny comparison between T2T C57BL/6J and CAST/EiJ shows the region is 3.45 Mbp in CAST/EiJ and harbours an inversion of 1.83 Mbp.

We found that the majority of filled gaps were consistent in size with the GRCm39 estimates (Supp Fig 1, Supplementary Fig. 1). However, we observe cases where gaps significantly expand the locus such as Chromosome 1 (GRCm39:85.32-85.3 Mbp, Figure 3C) where we have added an additional 3 Mbp compared to the gap of 49.9 Kbp. In both strains, this region is composed of highly repetitive sequences such as LINEs, SINEs and LTRs. We detected four inversions between C57BL/6J and CAST/EiJ, with inverted sequences exhibiting a notable difference in length between strains (685 Kbp in C57BL/6J versus 1,834 Kbp in CAST/EiJ, Supplementary Table 10). Additionally, we found that roughly 520 Kbp of sequence had been translocated in this locus in ten translocation events. Our analysis revealed 11 duplication events in C57BL/6J, totalling approximately 961 Kbp and 17 duplication events in CAST/EiJ, totalling approximately 572 Kbp. Our de novo gene annotation pipeline identified 55 and 59 protein-coding genes within this gap-filling region in C57BL/6J and CAST/EiJ respectively. The vast majority of these genes in both regions belong to the speckled protein gene family (C57BL/6J: 54.5%; CAST/EiJ: 59.3%, Supplementary Table 11). This gene family encode nuclear body proteins that are involved in innate and adaptive immune response and transcriptional regulation ^25^. We found all members of this gene family within this selected region (*Sp100, Sp110, Sp140, Sp140l2*) although individual gene counts differed between the strains. The most abundant speckled gene family member in this region was *Sp140l2* in both strains, with CAST/EiJ exhibiting four additional copies when compared to C57BL/6J (20 vs. 16). The largest gene count difference between C57BL/6J and CAST/EiJ was observed in *Sp140*, for which four additional copies were again found in CAST/EiJ (10 vs. 6). In contrast, C57BL/6J had three extra copies of *Sp100* that were not observed in this region in CAST/EiJ (5 vs. 3), whereas *Sp110* was found in equal counts in both strains (7). We observed five genes that were exclusively found in CAST/EiJ in this region. The most abundant was the zinc-finger protein *Cnbp* with two copies. The remaining four genes (*Rpl35, Hdlbp, Gm8113* and *BC035947*) exhibited a single copy within this region in CAST/EiJ. Only one gene was found exclusively in C57BL/6J in this region which was the zinc-finger protein gene *Zfp180*.

### Pseudoautosomal region

The pseudoautosomal region (PAR), which is shared by the X and Y chromosomes and located at the ends of them, contains numerous repeated sequences ^26^ and a high GC content ^27^, likely resulting from a high recombination frequency in this region (>100 times the genome average) ^28,29^one of the most challenging euchromatic regions to sequence and as a result only partial PAR sequences were included in GRCm39. In this study, we assembled the CAST/EiJ X chromosome PAR sequence, except for a large segmental duplication (SD) structure that could not be resolved (Figure 4A). We compared this to the C57BL/6J X chromosome PAR that was produced from diploid genome sequencing data and the PacBio walking method ^30^.

**Figure 4.**
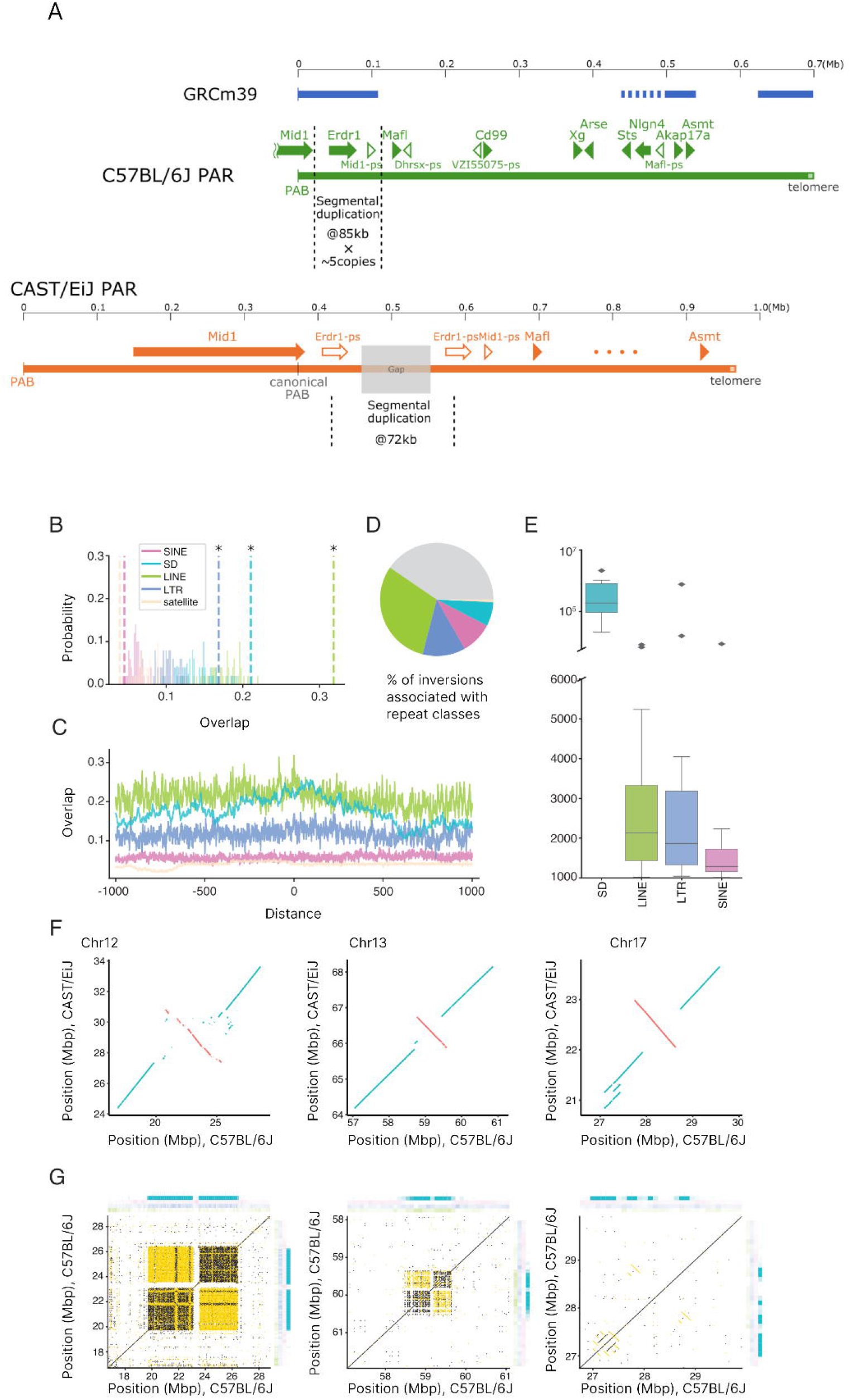
A) A comparison of the mouse PAR locus between GRCm39, C57BL/6J and T2T CAST/EiJ genomes. B) Expected overlap at inversion breakpoints from 1,000 random resampled permutations (histograms) compared to observed overlap (dotted lines) for different repeats. Asterisks denote significant enrichment (P<0.001). C) Repeat overlap (proportion of nucleotides attributed to a given repeat) calculated across 1 Kbp windows relative to inversion breakpoints in the GRCm39 genome (centred on zero) for various repeat types. D) Proportion of inversions associated with NAHR between different repeats. E) Length distributions for inversions associated with each considered genomic repeat. F) Dotplots for selected inversion regions generated from whole genome alignments between T2T C57BL/6J and CAST/EiJ. Collinear and inverted alignments are plotted in blue and orange, respectively. G) Dot plots generated from self-vs-self alignments of selected inversion breakpoint regions in the B6 genome. Collinear and inverted alignments are plotted in black and gold, respectively. Heatmaps show repeat density calculated across 10kb. Heatmap colours correspond to repeat colours used in (B-D).

In the mouse PAR, 10 genes (4 of which were novel), and 4 pseudogenes were identified (Figure 4A), which show synteny with the human PAR1. We located a large, tandem SD present between the *Mid1* and *Mafl* genes in both strains. The repeat unit size is approximately 85 Kbp in C57BL/6J and 72 Kbp in CAST/EiJ, and a copy number polymorphism is observed even in inbred mice. The copy number was quantified using digital PCR in 10 C57BL/6J mice purchased from the Jackson Laboratory, and a range of copy numbers per diploid was detected, with values between 2 and 23 (on average, 10.5). A comparison of the two strains revealed not only disparate locations of the PAR boundary, different sizes of the SD, and divergent physical sizes of the remaining region, but also a multitude of amino acid substitution mutations in all PAR genes (on average, one mutation per approximately 19 amino acid residues). The mouse PAR is a unique region of the mouse genome, exhibiting variation consistent with rapid change in sequence and structure.

### Inversions

Comparison of T2T assemblies revealed multiple large inversions between mouse strains and sheds light on their origins. Chromosomal inversions are genomic structural rearrangements in which a region of a chromosome is reversed between haplotypes. Inversions can play significant roles in evolution but are challenging to study since their breakpoints occupy highly repetitive genomic regions which are often poorly assembled ^31,32^. By comparing T2T assemblies, we identified 133 (>1kb) inversions between C57BL/6J and CAST/EiJ (Supplementary Table 12). Inversions are often formed through non-allelic homologous recombination (NAHR) between genomic repeats ^33,34^. Thus, to study the origins of inversions in house mice, we investigated repeat content at inversion breakpoints. We performed permutation tests comparing repeat overlap at inversion breakpoints to expectations from randomization as well as surrounding regions. Inversion breakpoints show enrichment for SDs as well as LINE and LTR retrotransposons, suggesting that these repeats may play roles in inversion formation (Figure 4B; permutation test, n=100 P<0.001 for all). LINE and LTR enrichment is highly localised and decays rapidly with distance from inversion breakpoints, consistent with the expected size of these TEs (∼200 bp - ∼8 kb) (Figure 4C). In contrast, SD enrichment extends hundreds of kilobases from inversion breakpoints, suggesting that inversions frequently inhabit complex SD-rich genomic regions (Figure 4C). To quantitatively estimate the number of inversions associated with different genomic repeats, we searched for patterns consistent with NAHR in which inversions are flanked by homologous repeats at both breakpoints ^34,35^. Overall, ∼60% of inversions show patterns consistent with NAHR, of these ∼50% are associated with LINEs, ∼21% with LTRs, ∼15% with SINEs, and ∼11% with SDs respectively (Figure 4D). Notably, although retrotransposons appear to have facilitated the majority of NAHR-mediated inversions SD-associated inversions are significantly longer (MWU, P<0.001; Figure 4E). Furthermore, larger SDs are associated with larger inversions (Kendall’s Tau = 0.63, P= 0.0007; Supplemental Figure 3). These results are consistent with observations in humans and deer mice suggesting that larger repeats are required to facilitate larger structural rearrangements ^35^. We then focused on the largest inversions, since longer inversions have more profound effects on genome structure and recombination. We identified several large inversions that are greater than 1 Mbp in length (Fig. 4F). Large inversions on chromosomes 12 and 13 both involve highly complex genomic regions primarily containing SDs (Fig. 4G). A large inversion on chromosome 17 shows long inverted repeats at its breakpoints, suggesting that it arose via NAHR (Fig. 4G).

### KRAB zinc finger loci

T2T assemblies have greatly improved the coverage of KRAB zinc finger proteins (KZFPs). KZFPs are one of the largest families of transcription factors in vertebrate genomes^1^. KZFPs preferentially bind transposable elements to recruit repressive epigenetic modifications ^36–38^. KZFPs are highly homologous and have been proposed to evolve through SDs, giving rise to a large number of KZFPs existing in clusters in mammalian genomes ^39,40^. These elements are highly polymorphic both between species ^36,37^ and between strains of mice, where strain-specific epigenetic modifiers have been identified in KZFP clusters ^41,42^. These loci are incomplete in GRCm39 limiting ability to effective profiling of the evolution and divergence of mouse KZFPs.

The distal arms of chromosomes 2 and 4 contain two of the largest clusters of KZFPs that are incomplete in GRCm39. The T2T assemblies have resolved the sequence of these KZFP clusters. For example, over 48 novel putative KZFPs in the C57BL/6J T2T genome have been identified. We observed large scale structural variations in KZFP clusters and significant divergence in the number of putative KZFPs between the two mouse strains (Fig. 5A and B). The chromosome 2 cluster in CAST/EiJ contains over 1.5 Mbp of additional sequence and an additional 42 putative KZFPs compared to the C57BL/6J genome (Fig. 5A). These additional genomic sequences have high homology to the elements present in the C57BL/6J genome, suggesting that they arose as the result of duplication events. The chromosome 4 KZFP cluster has a 2 Mbp inversion in the DNA sequence, as well as multiple potential SDs between the C57BL/6J and CAST/EiJ genome assemblies (Fig. 5B). However, other KZFPs clusters, such as the KZFP cluster on chromosome 17, do not show a significant improvement in sequence annotation using the T2T assemblies and do not show the same levels of polymorphism between mouse strains (Fig. 5C), indicating the distal KZFP clusters on chromosomes 2 and 4 are more highly variable and may be under additional evolutionary pressure.

**Figure 5.**
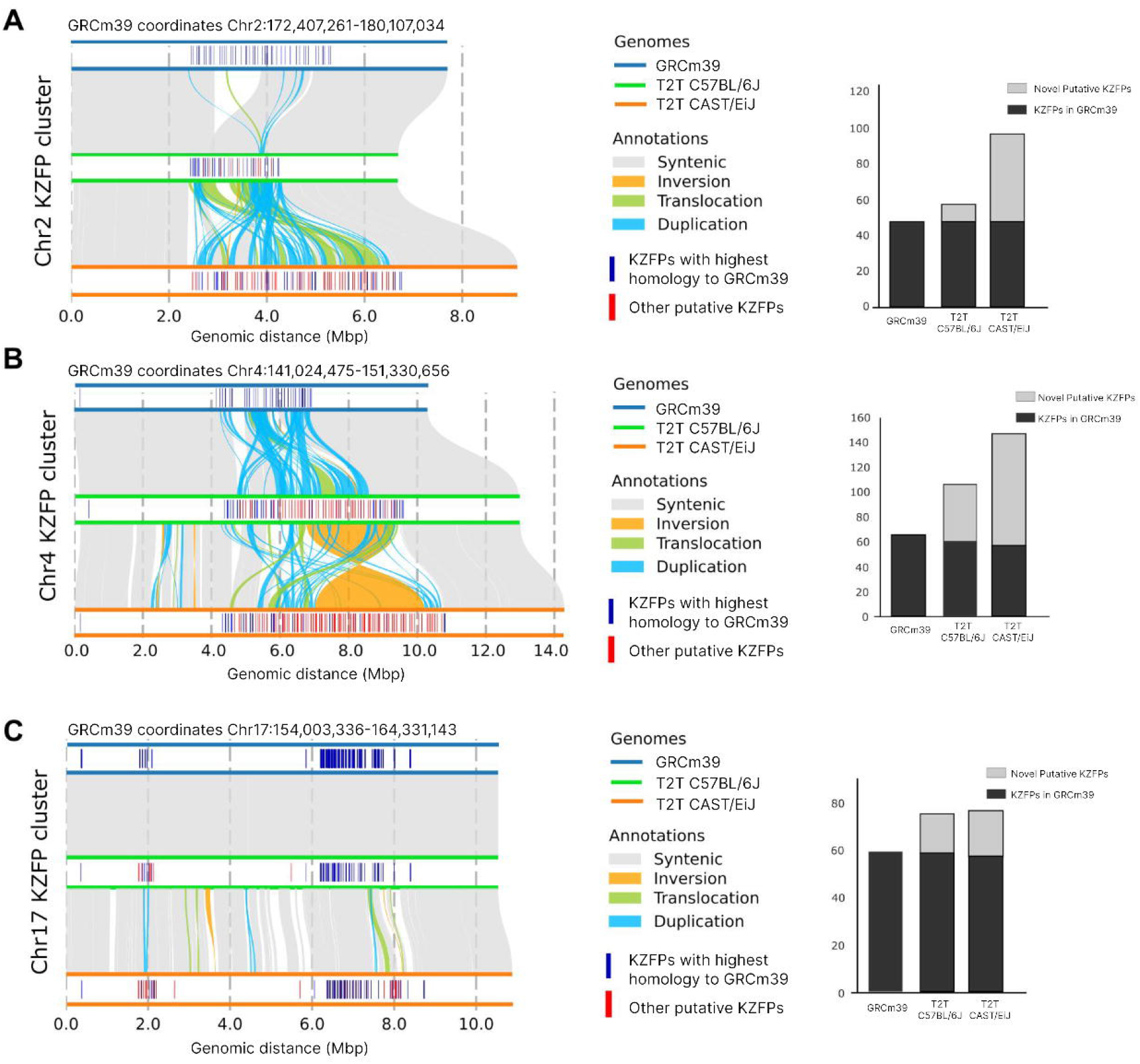
Comparison of the KRAB-zinc finger protein clusters between the GRCm39, T2T C57BL/6J, and CAST/EiJ genomes. Synteny plots for clusters on Chr2 (A), 4 (B), and 17 (C) are provided. Bar plots (right) give the numbers of KZFP proteins in each cluster per strain.

## Discussion

It is over twenty years since the first version of the mouse genome was released. The current version of the reference genome, GRCm39, remains incomplete and lacks sequence in many important loci. The work presented in this study represents a major milestone towards fully complete and accurate chromosomes for the mouse. We employed the latest ultra long sequencing technologies to produce T2T mouse reference genomes for two highly used strains (C57BL/6J and CAST/EiJ). We add 213 Mbp of sequence to the mouse reference genome containing an estimated 517 protein coding genes, providing sequence for all gaps in the current mouse reference. These genomes contain the first complete set of centromeres and telomeres for all autosomes and we carry out the first detailed comparative analysis of the mouse telomere and centromeres between two mouse subspecies revealing differences in both size and structure. Important loci such as the PAR locus, KZFP loci, and gap regions enriched for immunity loci can now be studied in much more detail as complete sequence will accelerate functional experiments and evolutionary analysis.

Inversions are copy neutral SVs that have the potential to disrupt the regulatory interactions of genes, and have proven the most challenging form of SVs to accurately detect using array and sequencing based approaches. We have shown how complete T2T genomes have made it possible to generate a complete set of inversions between the C57BL/6J and CAST/EiJ strains, and revealed several megabase scale inversions. The largest inversions are densely flanked by segmental duplications, a feature that has been identified in distant species such as deer mice and human, pointing towards a universal mechanism.

We highlighted a number of important loci which were missing or partially complete in GRCm39. For example, the KZFP clusters are known to be linked to strain-specific epigenetic outcomes, being able to annotate and properly profile the KZFPs across strains of mice is essential to our understanding of how the epigenetic landscape is established and evolves. From T2T genome assemblies, we are able to annotate and resolve previously un-resolved genomic elements, which are among the most divergent between mouse strains. This allows us to uncover the mechanisms behind how these elements evolve and drive divergent regulation of mammalian genomes.

This study represents a major milestone for mouse genetics that enables future functional studies in the incomplete regions of the mouse genome. Inbred and outbred hybrid mouse populations such as the Diversity Outbred Cross^43^ and Collaborative Cross^44^ are now being used to fine map a plethora of newly discovered QTL loci; the addition of complete sequence for key loci in two founder strains will accelerate this process. Future expansion of T2T reference genomes to further strains will form the basis for the Mouse Pangenome to fully represent mouse genetic diversity.

## Tables

Table 1. Genome statistics, quality metrics, repeats, and gene annotation

## Materials and Methods

See online materials and methods.

## Data Availability

The genome sequencing reads and assemblies are available from the European Nucleotide Archive under BioProject PRJEB47108, assembly accessions are GCA_964188535 (C57BL/6J) and GCA_964188545 (CAST/EiJ) (Supplementary Table 13). The genome assemblies and annotation are available via the Ensembl and the UCSC Genome Browsers.

## References

1. Little, C. C. A possible Mendelian explanation for a type of inheritance apparently non-Mendelian in nature. Science 40, 904–906 (1914).

2. Finlay, C. A., Hinds, P. W. & Levine, A. J. The p53 proto-oncogene can act as a suppressor of transformation. Cell 57, 1083–1093 (1989).

3. Takahashi, K. & Yamanaka, S. Induction of pluripotent stem cells from mouse embryonic and adult fibroblast cultures by defined factors. Cell 126, 663–676 (2006).

4. Mouse Genome Sequencing Consortium et al. Initial sequencing and comparative analysis of the mouse genome. Nature 420, 520–562 (2002).

5. Kipling, D., Ackford, H. E., Taylor, B. A. & Cooke, H. J. Mouse minor satellite DNA genetically maps to the centromere and is physically linked to the proximal telomere. Genomics 11, 235–241 (1991).

6. Joseph, A., Mitchell, A. R. & Miller, O. J. The organization of the mouse satellite DNA at centromeres. Exp. Cell Res. 183, 494–500 (1989).

7. Nurk, S. et al. The complete sequence of a human genome. Science 376, 44–53 (2022).

8. Rautiainen, M. et al. Telomere-to-telomere assembly of diploid chromosomes with Verkko. Nat. Biotechnol. 41, 1474–1482 (2023).

9. Cheng, H., Concepcion, G. T., Feng, X., Zhang, H. & Li, H. Haplotype-resolved de novo assembly using phased assembly graphs with hifiasm. Nat. Methods 18, 170–175 (2021).

10. Altemose, N. et al. Complete genomic and epigenetic maps of human centromeres. Science 376, eabl4178 (2022).

11. Sarsani, V. K. et al. The genome of C57BL/6J ‘Eve’, the mother of the laboratory mouse genome reference strain. G3 (Bethesda) 9, 1795–1805 (2019).

12. Kipling, D. & Cooke, H. J. Hypervariable ultra-long telomeres in mice. Nature 347, 400–402 (1990).

13. Prowse, K. R. & Greider, C. W. Developmental and tissue-specific regulation of mouse telomerase and telomere length. Proc. Natl. Acad. Sci. U. S. A. 92, 4818–4822 (1995).

14. Zhu, L. et al. Telomere length regulation in mice is linked to a novel chromosome locus. Proc. Natl. Acad. Sci. U. S. A. 95, 8648–8653 (1998).

15. Hemann, M. T. & Greider, C. W. Wild-derived inbred mouse strains have short telomeres. Nucleic Acids Res. 28, 4474–4478 (2000).

16. Kalitsis, P., Griffiths, B. & Choo, K. H. A. Mouse telocentric sequences reveal a high rate of homogenization and possible role in Robertsonian translocation. Proc. Natl. Acad. Sci. U. S. A. 103, 8786–8791 (2006).

17. Hörz, W. & Altenburger, W. Nucleotide sequence of mouse satellite DNA. Nucleic Acids Res. 9, 683–696 (1981).

18. Pardue, M. L. & Gall, J. G. Chromosomal localization of mouse satellite DNA. Science 168, 1356–1358 (1970).

19. Guenatri, M., Bailly, D., Maison, C. & Almouzni, G. Mouse centric and pericentric satellite repeats form distinct functional heterochromatin. J. Cell Biol. 166, 493–505 (2004).

20. Komissarov, A. S., Gavrilova, E. V., Demin, S. J., Ishov, A. M. & Podgornaya, O. I. Tandemly repeated DNA families in the mouse genome. BMC Genomics 12, 531 (2011).

21. Arora, U. P., Charlebois, C., Lawal, R. A. & Dumont, B. L. Population and subspecies diversity at mouse centromere satellites. BMC Genomics 22, 279 (2021).

22. Packiaraj, J. & Thakur, J. DNA satellite and chromatin organization at mouse centromeres and pericentromeres. Genome Biol. 25, 52 (2024).

23. Church, D. M. et al. Lineage-specific biology revealed by a finished genome assembly of the mouse. PLoS Biol. 7, e1000112 (2009).

24. Lilue, J. et al. Sixteen diverse laboratory mouse reference genomes define strain-specific haplotypes and novel functional loci. Nat. Genet. 50, 1574–1583 (2018).

25. Fraschilla, I. & Jeffrey, K. L. The Speckled Protein (SP) family: Immunity’s chromatin readers. Trends Immunol. 41, 572–585 (2020).

26. Takahashi, Y. et al. Methylation imprinting was observed of mouse mo-2 macrosatellite on the pseudoautosomal region but not on chromosome 9. Chromosoma 103, 450–458 (1994).

27. Kasahara, T., Abe, K., Mekada, K., Yoshiki, A. & Kato, T. Genetic variation of melatonin productivity in laboratory mice under domestication. Proc. Natl. Acad. Sci. U. S. A. 107, 6412–6417 (2010).

28. Kipling, D., Salido, E. C., Shapiro, L. J. & Cooke, H. J. High frequency de novo alterations in the long-range genomic structure of the mouse pseudoautosomal region. Nat. Genet. 13, 78–80 (1996).

29. Lange, J. et al. The landscape of mouse meiotic double-strand break formation, processing, and repair. Cell 167, 695–708.e16 (2016).

30. Kasahara, T., Mekada, K., Abe, K., Ashworth, A. & Kato, T. Complete sequencing of the mouse pseudoautosomal region, the most rapidly evolving ‘chromosome’. Genomics (2022).

31. Wellenreuther, M. & Bernatchez, L. Eco-evolutionary genomics of chromosomal inversions. Trends Ecol. Evol. 33, 427–440 (2018).

32. Mérot, C., Oomen, R. A., Tigano, A. & Wellenreuther, M. A roadmap for understanding the evolutionary significance of structural genomic variation. Trends Ecol. Evol. 35, 561–572 (2020).

33. Cáceres, M., Ranz, J. M., Barbadilla, A., Long, M. & Ruiz, A. Generation of a widespread Drosophila inversion by a transposable element. Science 285, 415–418 (1999).

34. Porubsky, D. et al. Recurrent inversion toggling and great ape genome evolution. Nat. Genet. 52, 849–858 (2020).

35. Porubsky, D. et al. Recurrent inversion polymorphisms in humans associate with genetic instability and genomic disorders. Cell 185, 1986–2005.e26 (2022).

36. Imbeault, M.Helleboid, P.-Y. & Trono, D. KRAB zinc-finger proteins contribute to the evolution of gene regulatory networks. Nature 543, 550–554 (2017).

37. de Tribolet-Hardy, J. et al. Genetic features and genomic targets of human KRAB-zinc finger proteins. Genome Res. 33, 1409–1423 (2023).

38. Shi, H. et al. ZFP57 regulation of transposable elements and gene expression within and beyond imprinted domains. Epigenetics Chromatin 12, 49 (2019).

39. Huntley, S. et al. A comprehensive catalog of human KRAB-associated zinc finger genes: insights into the evolutionary history of a large family of transcriptional repressors. Genome Res. 16, 669–677 (2006).

40. Kauzlaric, A. et al. The mouse genome displays highly dynamic populations of KRAB-zinc finger protein genes and related genetic units. PLoS One 12, e0173746 (2017).

41. Ratnam, S. et al. Identification of Ssm1b, a novel modifier of DNA methylation, and its expression during mouse embryogenesis. Development 141, 2024–2034 (2014).

42. Bertozzi, T. M., Elmer, J. L., Macfarlan, T. S. & Ferguson-Smith, A. C. KRAB zinc finger protein diversification drives mammalian interindividual methylation variability. Proc. Natl. Acad. Sci. U. S. A. 117, 31290–31300 (2020).

43. Churchill, G. A., Gatti, D. M., Munger, S. C. & Svenson, K. L. The Diversity Outbred mouse population. Mamm. Genome 23, 713–718 (2012).

44. Collaborative Cross Consortium. The genome architecture of the Collaborative Cross mouse genetic reference population. Genetics 190, 389–401 (2012).

