## Supplementary material for "The structural diversity of telomeres and centromeres across mouse subspecies revealed by complete assemblies": Methods

### Materials and Methods

**Mice:** Mouse embryonic stem cells were derived from 3.5 dpc F1 embryos resulting from a cross between CAST/EiJ (RRID:IMSR_JAX:000928) dams and C57BL/6J (RRID:IMSR_JAX:000664) sires. Derivation and characterization (mycoplasma testing, SNP genotyping, pluripotency marker expression, chromosome counting, germline testing) of the mESCs were as previously described [1]. One male line (CASTB6-9) with a >70% euploid karyotype was selected for sequencing. CASTB6-9 was cultured as previously described [1], dissociated, washed in PBS, and then pelleted before being flash-frozen in liquid nitrogen and stored at -80°C. $5x10^6$ and $1x10^6$ cell aliquots were shipped on dry ice to the Sanger Institute for sequencing. For the digital PCR measurement of PAR regions, C57BL/6J mice (RRID:IMSR_JAX:000664) were used. All procedures involving laboratory mice were approved by The Institutional Animal Care and Use Committees of The Jackson Laboratory (under Animal Use Summary #20030) and RIKEN.

**DNA and sequencing:** For PacBio HiFi and ONT ultra-long sequencing, high molecular weight (HMW) DNA was extracted from 1M (PacBio) and 5M (ONT) cell pellets using the Monarch T3050 kit and protocol (New England BioLabs). For ONT ultra-long, recommended protocol adjustments were carried out. DNA QC was performed using FemtoPulse, Qubit measurement taken from the top, middle and bottom of the extractions, and homogenisation of extracted DNA by gently pipetting with a wide bore pipette tip. Qubit measurements from the top, middle, and bottom of the tube were repeated until values were similar. Library preparation for PacBio sequencing using template preparation kit 2.0. Sequencing on PacBio Sequel IIe with SMRT Cell 8M using Binding Kit 2.2, and Sequencing kit 2.0a, to achieve 50X coverage (6 SMRT Cells were run in total). ONT Library preparation was performed with ONT's UL sequencing kit (ULK001), followed by sequencing on PromethION 10.4.1 (runs were monitored to perform multiple nuclease flushes and reload more library to maximise yield from each flow cell).

**Genome assembly:** Our C57BL/6J and CAST/EiJ assemblies were generated using a combination of haplotype-aware, T2T-capable genome assembly approaches: (i) Verkko (v1.3.1); and (ii) Hifiasm (v0.19.5). For each assembler, we generated multiple assemblies using different combinations of read quality and read length subsets. All assemblies were executed in trio-binning mode using both the HiFi and ONT reads and k-mer databases generated from strain-specific Illumina short reads as input. The parental k-mer databases were generated by Merqury (v1.3) [2] and Yak (v0.1-r66-dirty, https://github.com/lh3/yak) for Verkko and Hifiasm respectively, following each assembler's recommended guidelines. For each of our assembly runs, this method produced a set of haplotype-separated contigs for each strain which were then ordered and oriented into chromosome scaffolds with RagTag (v2.1.0) [3], using each strain's respective reference genome (C57BL/6J: GCA_000001635.9; CAST/EiJ: GCA_921999005.2) as an anchor.

**Assembly Evaluation:** All assemblies were evaluated and compared to one another to identify the best set of chromosome scaffolds to serve as each strain's respective base genome assembly. Merqury (v1.3) was used to assess the k-mer completeness of our assemblies. Haplotype separation refers to the process of separating a diploid assembly into each of its haploid components. In the context of the mouse T2T assemblies, this refers to separating our diploid F1 sample into a C57BL/6J assembly and a CAST/EiJ assembly. Haplotype separation in each of our assemblies was evaluated using the "trioeval" command in Yak (v0.1-r66-dirty, https://github.com/lh3/yak) which compares the k-mer spectrum from each haplotype-separated assembly to k-mer spectrums generated from Illumina reads from each parental strain. These comparisons are used to calculate switch error and hamming error rates which are quantitative measures of haplotype separation. Haplotype separation was further evaluated by aligning haplotype-separated contigs to a combined GRCm39 and a pure CAST/EiJ long read reference genome [GCA_921999005.2] with minimap2 (v2.24-r1122) [4]. Mouse canonical telomeric repeats (TTAGGGn) were detected in each of our assemblies using the "telo" command in seqtk (https://github.com/lh3/seqtk) which searches for telomeric repeats at the end of each sequence within a FASTA file to evaluate if our newly generated mouse chromosomes properly terminated in canonical telomeric repeat. Finally, we also compared the number of hybrid structural variant calls reported by Sniffles (v2.0.7) [5] using both HiFi and ONT read alignment and used these counts as a measure of assembly accuracy.

We used the information generated above to compare and rank each of our candidate mouse strain assemblies against each other. We chose the assembly that ranked the highest across our chosen criteria to become our base C57BL/6J and CAST/EiJ chromosomes (Supplementary Table 1). Both selected assemblies for each strain were generated using Verkko. For each strain, chromosome-to-chromosome alignment comparisons were generated between our assemblies with Winnowmap (v2.03) [6]. When we identified cases where a given telomeric region was missing in the base assembly chromosomes but was present in a given secondary assembly, we incorporated the corresponding sequences from the secondary assembly into the base chromosomal assembly. All assembly changes were supported by both HiFi and ONT read alignments generated by Winnowmap where multiple reads spanned our integration boundary.

**Assembly Polishing:** We performed a hybrid HiFi and ONT read-based error correction pipeline, previously outlined in [7], to correct any remaining structural errors in our C57BL/6J and CAST/EiJ assemblies. Briefly, Sniffles was used to call structural variants (SVs) in our assemblies using both HiFi and ONT read alignments (v2.0.7)[5], and insertion and deletion sequences from these SV calls were then polished using Iris (v1.0.4) (https://github.com/mkirsche/Iris). Next, Jasmine (v1.1.5) (https://doi.org/10.1101/2021.05.27.445886) was used to merge our independent HiFi and ONT call sets to identify all variants that were observed using both sequencing technologies. Finally, we filtered these shared variants using Merfin [8] and incorporated these SV corrections into our final assemblies with bcftools [9]. This polishing process improved the base accuracy of both assemblies (C57BL/6J: 47.7 to 54.9 QV; CAST/EiJ: 44.4 to 44.6 QV).

**Assembly Curation:** The T2T genomes were manually curated using Hi-C data. In brief, Hi-C reads were mapped to the T2T genomes following the Arima Hi-C mapping pipeline (https://github.com/ArimaGenomics/mapping_pipeline). We then generated and visualised a Hi-C contact map using PretextMap and PretextView which was used to manually curate our chromosomes. Following this process, several of our mouse chromosomes still did not end in telomeric sequence on their centromeric end. We identified these 'missing' telomere sequences by searching for the mouse canonical telomere repeat in our assemblies' unplaced contigs. We noticed that many of these contigs also contained centromere repeats, supporting their placement in the 'missing' regions in our assemblies. To assign these telocentric contigs to the correct chromosome, we employed a mapping-based approach using MashMap (v3.1.1) [10]. It has previously been noted in human studies that large satellite arrays tend to have more similarity within a given chromosome array than between different chromosomes [11]. Therefore, we used MashMap to map our unplaced telocentric sequences against our chromosome-assigned scaffolds and quantified their sequence similarity to each chromosome scaffold, which we then used to identify the chromosome scaffold with the highest amount of similarity. To support this approach, we also quantified the amount of Hi-C read pairs with one mate on a given unplaced telocentric sequence and a given chromosome scaffold to provide supporting evidence of linkage to a particular chromosome. Using information from these approaches, we assigned all remaining telocentric sequences to a chromosome during which we introduced a model gap of 100bp between the contig and the chromosome. This result of this process now meant that all of our mouse chromosomes ended in mouse canonical telomere repeat on both ends. The final chromosomes are available under accessions GCA_964188545.1 (CAST/EiJ) and GCA_964188535.1 (C57BL/6J).

**Gene prediction and annotation:** We used BRAKER3 (v3.0.3) [12] to predict protein-coding gene structures in our assemblies using both RNA-seq and protein evidence to train the gene prediction pipeline. RNA-seq data was acquired from the ENCODE portal [13] and public databases (Supplementary Table 4). RNA-seq reads were then aligned to each strain's respective genome using STAR (v2.7.10b) [14]. For protein evidence, we used the Vertebrata database acquired from OrthoDB[15].

In addition to our BRAKER3 de novo gene prediction, we also produced an annotation transferring genes from GRCm39 to our new T2T assemblies using Liftoff [16] using the following arguments: -copies -sc 0.95 -polish -exclude_partial.

**Repetitive Sequence Annotation:** We used RepeatMasker (v4.1.5) (http://www.repeatmasker.org) with the default Dfam repetitive element library in "mus musculus" mode to identify and annotate repetitive elements in our new C57BL/6J and CAST/EiJ genomes. To validate and refine our repeat annotations, we further supported our RepeatMasker annotations with targeted BLAST searches using C57BL/6J reference sequences for the minor satellite and TLC repeats [17–20].

**Identification of novel genes**

To identify potential novel genes in the T2T genomes, we extracted genes from each BRAKER annotation that exhibited no overlap with any gene from the Liftoff annotation using bcftools intersect -v. The output GFF3 file was then filtered to include only BRAKER gene entries that had >= 3 exons and >= 200 base pairs of coding sequence. The protein sequences for these filtered genes were then used as query sequences for a BLASTp search against all C57BL/6J proteins in the Ensembl genome browser[21]. Each BRAKER gene was assigned a top BLAST hit from this search to infer its potential function (Supplemental Table 5).

**PANTHER protein classification**

We used the PANTHER (Protein ANalysis THrough Evolutionary Relationships) classification system (v19.0)[22] to assign protein classes to genes of interest within our Liftoff annotations (Supplementary Table 6). For a given Liftoff gene, this was achieved by inputting its' associated Ensembl gene ID, lifted over from GRCm39 annotation, into the PANTHER web-based server.

**Identification of gap-filling sequences:** To identify the sequences in our new C57BL/6J assembly that fill the assembly gaps observed in GRCm39, we implemented an alignment-based approach using the repeat-sensitive alignment software Winnowmap (v2.03) [6]. Firstly, we extracted the flanking sequences of all 87 gaps in GRCm39 autosomes and aligned them to our new C57BL/6J assembly with Winnowmap. Gap-filling sequences were then inferred as the sequence between each gap's left and right flanking sequence alignments. Using these gap-filling sequences, we characterised gaps as: completely filled (novel sequence added with no gap bases remaining); partially filled (novel sequence added with some gap bases still remaining); or not filled (only one/no flanking sequence alignment or no non-N bases added).

**PAR assembly using PacBio walking method:** PacBio HiFi reads from C57BL/6 J genomic DNA (SRR11606870) [23] were mapped to known "seed" sequences, *i.e.*, any of the exons of *Asmt*, *Akap17a*, *Mafl-ps*, *Nlgn4*, *Sts*, *Arse*, *Mafl*, *Erdr1*, *Mid1*, and *Gm52481* genes, using minimap2 (ver. 2.17-r941) with the parameters -k 27 -w 18 -m 99. The alignments were processed using samtools (ver. 1.1) and visualised with IGV (ver. 2.8.13). We manually selected reads that were identical (except for obvious mutations or polymorphisms in the seed sequence) to the seed sequence over 4 kb and assembled them using CAP3 (VersionDate, 12/21/07) with the default parameters. If more than two contigs were generated, the contig consisting of the largest number of reads was used as the representative. To visualise the hallmark of the contig sequence, we created a dot-plot view of self-similarity using web YASS (https://bioinfo.lifl.fr/yass/yass.php) or local YASS (ver. 1.15) with the default parameters. We compared the self-similarity view of the contig with that of the seed sequence and confirmed that the walking was proceeding correctly. When the self-similarity view of the contig that we took as representative was obviously different from that of the seed sequence, we used another contig as an alternative representative. Next, the representative contig and the seed sequence were compared using BLASTN 2 sequences program (default parameters) and manually merged them. Basically, the contig sequence was connected to the seed sequence near the centre where these sequences overlapped. A 10-kb sequence from the end of the merged sequence was used as the new seed for the next round of walking. Each round of walking yielded a new sequence of 3.5-12.5 kb (~9.8 kb on average).

**PAR Digital PCR:** Genome DNA was extracted from liver, brain, or tail chip of C57BL/6J mice using Monarch Genomic DNA Purification Kit (Cat# T3010S, New England Biolabs) and digested by *Pst* I (Cat# R3140S, New England Biolabs). After heat inactivation (60 °C for 15 min) and dilution with water (5 ng/μL), we performed digital PCR using the QuantStudio 3D system (Thermo Fisher Scientific) according to the manufacturer's procedure. We used TaqMan Copy Number Reference Assay, mouse, Tfrc (Cat# 4458366) to count the chromosome 16 (2 copies per the diploid genome) and Custom TaqMan Copy Number Assays (mMid1Ex5 and mMid1Ex7), which targeted the exons 5 and 7 of *Mid1* gene, designed by the TaqMan Custom Design Assay Tool (Thermo Fisher Scientific). The sequences of the primers and probes are as follows:

mMid1Ex5: CTCAGCAGATTGCAAACTGTAAACA, GGCGTGGTCATTTTCCTTCAG, and [FAM-]TCTGCATCGCTCATCTCGC
mMid1Ex7: CTGCACCGCTTCCTACGA, GTAGGAGACCACGCTGAACTC, and [FAM-]CCACTGGACCTCAGAGGAC

The copy numbers of the SD obtained by using mMid1Ex5 and mMid1Ex7 assays were the same. The copy numbers in DNA samples extracted from the liver, brain, and tail tip of the same individual were the same, indicating that the copy number did not change during ontogeny.

**Inversions Methods:** To identify inversions between C57BL/6J and CAST/EiJ strains, we first aligned C57BL/6J and CAST/EiJ T2T genomes using minimap2 (version 2.21) with flags -a --eqx -x asm5 --cs -r2k [4]. We sorted and indexed resulting bam files using samtools (version 1.10) [24] and called inversions using SyRI (version 1.6.3) [25], which uses alignment of syntenic regions to accurately detect structural rearrangements. SyRI performs particularly well in identifying large balanced structural variants such as inversions from whole genome alignments compared to other available tools. Due to the challenges with systematically calling balanced structural variants in highly repetitive regions, we filtered out inversions primarily composed of simple repeats and satellites and manually inspected dotplots to filter out spurious calls [26–28]. We also filtered out erroneous balanced inversion calls likely caused by twin-priming during L1 retrotransposition by removing inversions covered ≥95% by L1 models [29]. After filtering, we were left with 131 inversions larger than 1 kb (Supplemental Table 12).

To investigate the genomic mechanisms underlying inversions in house mice, we explored repeats at inversion breakpoint regions. We first performed permutation tests for enrichment of repeats in inversion breakpoint regions for five types of repeats: LINEs, SINEs, LTR retrotransposons, satellites and segmental duplications. Specifically, we obtained the 1 kb flanking regions surrounding each inversion breakpoint using bedtools flank (version 2.29.1)[30]. We then assessed various metrics of repeat composition in these regions, comparing them to expectations derived from 1,000 randomly resampled permutations using GAT (version 1.3.5) [31]. Considered metrics included the count of repeats intersecting with the inversion breakpoint regions and the percentage of base pairs in breakpoint regions associated with specific repeats. To search for evidence of repeat-mediated inversions, we intersected inversion breakpoint regions with TE and SD annotations. Using the 500 bp regions flanking each inversion, we called repeat-mediated inversions based on the presence of TEs from the same family at both breakpoints, or flanking SDs at both breakpoints. To investigate the relationship between inversion length and associated SD length, we performed a linear regression comparing inversion length to mean adjacent SD length, finding a significant correlation (Kendall's Tau = 0.63, P= 0.0007). To visualize SD-enriched inversion breakpoints, we generated self-vs-self alignments of breakpoint regions using minimap2 with the flag -P and produced dotplots using python.

### References

1. Czechanski, A. *et al.* Derivation and characterization of mouse embryonic stem cells from permissive and nonpermissive strains. *Nat. Protoc.* **9**, 559–574 (2014).
2. Rhie, A., Walenz, B. P., Koren, S. & Phillippy, A. M. Merqury: reference-free quality, completeness, and phasing assessment for genome assemblies. *Genome Biol.* **21**, 245 (2020).
3. Alonge, M. *et al.* Automated assembly scaffolding using RagTag elevates a new tomato system for high-throughput genome editing. *Genome Biol.* **23**, 258 (2022).
4. Li, H. Minimap2: pairwise alignment for nucleotide sequences. *Bioinformatics* **34**, 3094–3100 (2018).
5. Sedlazeck, F. J. *et al.* Accurate detection of complex structural variations using single-molecule sequencing. *Nat. Methods* **15**, 461–468 (2018).
6. Jain, C. *et al.* Weighted minimizer sampling improves long read mapping. *Bioinformatics* **36**, i111–i118 (2020).
7. Mc Cartney, A. M. *et al.* Chasing perfection: validation and polishing strategies for telomere-to-telomere genome assemblies. *Nat. Methods* **19**, 687–695 (2022).
8. Formenti, G. *et al.* Merfin: improved variant filtering, assembly evaluation and polishing via k-mer validation. *Nat. Methods* **19**, 696–704 (2022).
9. Danecek, P. *et al.* Twelve years of SAMtools and BCFtools. *Gigascience* **10**, (2021).
10. Kille, B., Garrison, E., Treangen, T. J. & Phillippy, A. M. Minmers are a generalization of minimizers that enable unbiased local Jaccard estimation. *Bioinformatics* **39**, (2023).
11. Altemose, N. *et al.* Complete genomic and epigenetic maps of human centromeres. *Science* **376**, eabl4178 (2022).
12. Gabriel, L. *et al.* BRAKER3: Fully automated genome annotation using RNA-seq and protein evidence with GeneMark-ETP, AUGUSTUS, and TSEBRA. *Genome Res.* **34**, 769–777 (2024).
13. Luo, Y. *et al.* New developments on the Encyclopedia of DNA Elements (ENCODE) data portal. *Nucleic Acids Res.* **48**, D882–D889 (2020).
14. Dobin, A. *et al.* STAR: ultrafast universal RNA-seq aligner. *Bioinformatics* **29**, 15–21

(2013).

15. Kuznetsov, D. *et al.* OrthoDB v11: annotation of orthologs in the widest sampling of organismal diversity. *Nucleic Acids Res.* **51**, D445–D451 (2023).
16. Shumate, A. & Salzberg, S. L. Liftoff: accurate mapping of gene annotations. *Bioinformatics* **37**, 1639–1643 (2021).
17. Vissel, B. & Choo, K. H. Mouse major (gamma) satellite DNA is highly conserved and organized into extremely long tandem arrays: implications for recombination between nonhomologous chromosomes. *Genomics* **5**, 407–414 (1989).
18. Radic, M. Z., Lundgren, K. & Hamkalo, B. A. Curvature of mouse satellite DNA and condensation of heterochromatin. *Cell* **50**, 1101–1108 (1987).
19. Kalitsis, P., Griffiths, B. & Choo, K. H. A. Mouse telocentric sequences reveal a high rate of homogenization and possible role in Robertsonian translocation. *Proc. Natl. Acad. Sci. U. S. A.* **103**, 8786–8791 (2006).
20. Kipling, D., Wilson, H. E., Mitchell, A. R., Taylor, B. A. & Cooke, H. J. Mouse centromere mapping using oligonucleotide probes that detect variants of the minor satellite. *Chromosoma* **103**, 46–55 (1994).
21. Harrison, P. W. *et al.* Ensembl 2024. *Nucleic Acids Res.* **52**, D891–D899 (2024).
22. Thomas, P. D. *et al.* PANTHER: Making genome-scale phylogenetics accessible to all. *Protein Sci.* **31**, 8–22 (2022).
23. Hon, T. *et al.* Highly accurate long-read HiFi sequencing data for five complex genomes. *Sci. Data* **7**, 399 (2020).
24. Li, H. *et al.* The Sequence Alignment/Map format and SAMtools. *Bioinformatics* **25**, 2078–2079 (2009).
25. Goel, M., Sun, H., Jiao, W.-B. & Schneeberger, K. SyRI: finding genomic rearrangements and local sequence differences from whole-genome assemblies. *Genome Biol.* **20**, 277 (2019).
26. Mahmoud, M. *et al.* Structural variant calling: the long and the short of it. *Genome Biol.* **20**, 246 (2019).

27. Porubsky, D. *et al.* Inversion polymorphism in a complete human genome assembly. *Genome Biol.* **24**, 100 (2023).

28. Porubsky, D. *et al.* Recurrent inversion polymorphisms in humans associate with genetic instability and genomic disorders. *Cell* **185**, 1986–2005.e26 (2022).

29. Ostertag, E. M. & Kazazian, H. H., Jr. Twin priming: a proposed mechanism for the creation of inversions in L1 retrotransposition. *Genome Res.* **11**, 2059–2065 (2001).

30. Quinlan, A. R. & Hall, I. M. BEDTools: a flexible suite of utilities for comparing genomic features. *Bioinformatics* **26**, 841–842 (2010).

31. Heger, A., Webber, C., Goodson, M., Ponting, C. P. & Lunter, G. GAT: a simulation framework for testing the association of genomic intervals. *Bioinformatics* **29**, 2046–2048 (2013).
