## Supplementary material for "The structural diversity of telomeres and centromeres across mouse subspecies revealed by complete assemblies": Table 1

| | GRCm39 | T2T-C57BL/6J | T2T-CAST/EiJ |
|---|---|---|---|
| **Assembly** | | | |
| Total ungapped length, autosomes (Gbp) | 2.397 | 2.638 | 2.665 |
| Assembly QV | 54.42 | 54.93 | 48.24 |
| Canonical telomeres | 6 | 38 | 38 |
| Canonical telomere pairs | 0 | 19 | 19 |
| **Annotation** | | | |
| Protein-coding genes (BRAKER) | | 21,423 | 21,440 |
| Protein-coding genes (LiftOff) | | 21,469 | 21,490 |
| **Repetitive Bases (Mbp)** | | | |
| SINEs | 158.954 | 160.028 | 160.736 |
| LINEs | 445.782 | 450.197 | 438.161 |
| LTR elements | 262.932 | 267.257 | 266.510 |
| DNA elements | 19.473 | 19.551 | 19.614 |
| Unclassified | 13.295 | 13.381 | 13.711 |
| Interspersed repeats | 900.437 | 910.416 | 898.734 |
| Small RNA | 1.304 | 1.365 | 1.425 |
| Satellites | 6.659 | 189.983 | 227.884 |
| Simple repeats | 61.688 | 64.506 | 67.900 |
| Low complexity | 8.677 | 9.002 | 8.956 |
| Total repetitve bases | 1879.201 | 2085.686 | 2103.631 |
